# ZymePackNet: rotamer-sampling free graph neural network method for protein sidechain prediction

**DOI:** 10.1101/2023.05.05.539648

**Authors:** Abhishek Mukhopadhyay, Amit Kadan, Benjamin McMaster, J. Liam McWhirter, Surjit B. Dixit

## Abstract

Protein sidechain conformation prediction, or packing, is a key step in many in silico protein modeling and design tasks. Popular protein packing methods typically rely on approximated energy functions and complex algorithms to search dense rotamer libraries. Inspired by the recent success of deep learning in protein modeling tasks, we present ZymePackNet, a graph neural network based protein packing tool that does not require a rotamer library, scoring functions or a search algorithm. We train regression models using protein crystal structures represented as graphs, which are employed sequentially to “germinate” the sidechain starting from atoms anchoring the protein backbone to the sidechains’ termini, followed by an iterative refinement stage. ZymePackNet is fast and accurate compared to state-of-the-art protein packing methods. We validate our model on three native backbone datasets achieving a mean average error of 16.6°, 24.1°, 42.1°, and 53.0° for sidechain dihedral angles (*χ*_1_ to *χ*_4_). ZymePackNet captures complex physical interactions such as *π* stacking without explicitly accounting for it in the model; such effects are currently lacking in the energy terms used in traditional packing tools.

**Supplementary information:** Supplementary data are available at *Bioinformatics* online.

## 1 Introduction

Accurate prediction of sidechain conformation is a key component of protein modelling, mutagenesis, protein structure prediction, and protein engineering and design problems. In single-site mutants and in homologous proteins, the backbone conformation change may be minimal and structure prediction can often be accomplished by accurately predicting the sidechain conformation of the mutated region only. Sidechain geometry is also key for binding site recognition, for *in silico* binding affinity assessment, and for interface engineering between cognate binding proteins and protein/ligand complexes. For structure refinement methods that include backbone conformation change, one stage in the refinement process is prediction of side-chain conformation, also known as repacking. While accuracy is important, speed is key for refining a large ensemble of decoys with different backbone geometry using *in silico* methods. For the protein design problem, one needs to co-optimize changes in the sequence along with backbone and side chain conformations; consequently, in the absence of expert or prior knowledge about the couplings between the mutation sites, there can be a combinatorial explosion of possible solutions.

There is a plethora of sidechain packing algorithms that successfully attempt to solve the sidechain packing problem to varied degrees of accuracy or computational efficiency. Traditional packing methods (Simonson *et al*., 2013; Miao *et al*., 2011; Cao *et al*., 2011; Huang *et al*., 2020; Liang *et al*., 2011; Krivov *et al*., 2009) comprise broadly of three components: (i) a rotamer library, (ii) an energy/scoring function, and (iii) a search algorithm. These methods use search algorithms to predict a set of rotamers, one rotamer for each amino acid, for the region of interest in a protein to minimize the overall energy of the system. Rotamer libraries (Dunbrack Jr, 2002; Xiang and Honig, 2001; Shapovalov and Dunbrack Jr, 2011) are a collection of frequencies, mean dihedral angles, and standard deviations of the discrete conformations (rotamers) of the amino acid sidechains derived from proteins in the crystal PDB database. Broadly, there are two categories of rotamer libraries – a backbone-dependent rotamer library (BBDRL), where the frequencies, mean dihedral angles, and standard deviations of the rotamers are a function of the protein backbone dihedral angles, and backbone-independent rotamer libraries (BBIRL), where the frequencies and mean dihedral angles are independent of the backbone conformation. Compared to BBRIL, BBDRL is preferred by most modern packing algorithms because of its smaller conformational search space (Huang *et al*., 2019). Nevertheless, even BBDRLs are generic and lack details of residues’ local environment, resulting in a computational burden incurred for scouting parts of the rotamer space that are highly energetically unfavorable. Certain terms in the scoring function are quite standard and commonplace amongst available methods; these terms (Miao *et al*., 2011; Cao et al., 2011; Huang *et al*., 2020; Krivov *et al*., 2009) account for Van Der Waals interactions, hydrogen bonding, disulfide bonding, and rotamer frequency. Due to analytical complexity, scoring functions are typically overly simplified or intricately parameterized to maintain reasonable accuracy while being computationally efficient. For example, non-covalent interactions energy terms, such as the ones that account for electrostatics, solvation effects (Pokala and Handel, 2004; Zhu et al., 2006; Simonson *et al*., 2013; Mukhopadhyay et al., 2015; Onufriev and Case, 2019) or aromatic *π* stacking (Li *et al*., 2011), are empirical or semi-empirical models that can be too expensive to evaluate even with tractable numerical methods. For sampling, while exhaustive enumeration is feasible to pack a smaller number of amino acid sites or a conformation space limited by a sparse rotamer library, for most real-world problems it is not tractable as the number of possible solutions scales exponentially. Sampling algorithms can be grouped into two types: deterministic algorithms that guarantee finding a global or at least a local minimum, e.g. Dead-End Elimination (Desmet *et al*., 1992), Iterated Conditional Modes (Besag, 1986; Xiang and Honig, 2001), Integer Linear Programming(Zhu, 2007; Kingsford *et al*., 2005; Saraf et al., 2006), Branch and Bound (Gordon and Mayo, 1999; Traoré *et al*., 2016), Tree Decomposition (Xu *et al*., 2005) and stochastic or heuristic algorithms, e.g. Metropolis (Samish, 2017), Parallel Tempering (Yang and Saven, 2005; Druart *et al*., 2016), Simulated Annealing (Lee and Subbiah, 1991; Leaver-Fay *et al*., 2011), Genetic Algorithms (Jones, 1994). Even if their implementation is parallelized, such sampling algorithms are also limited by the exponential scaling of the solution space, losing any sizable advantage gained from the parallelization.

Recently, deep learning has gained popularity in this field due to its demonstrated ability in effectively leveraging sequence and structure data available in the public domain to solve difficult problems like protein modeling and design. Some recent methods have also emerged that harness applied machine learning methods for sidechain modeling. Most of these methods are discriminative models that score rotamers given the protein environment about a single amino acid site (Du *et al*., 2020) or replace physics-based energy with neural network derived probabilistic scores for every rotamer in the conformational space defined by rotamer libraries (Liu *et al*., 2017; Misiura et al., 2021). Another recent method by Xu *et al*. (2020) uses a hybrid approach where they enrich the rotamer library with neural network derived rotamers and combine results of various sidechain prediction results via an ensemble method to sample the conformational space using simple energy terms e.g., Van der Waals like pair energy terms and a rotamer-frequency based energy term.

Interestingly, inspired by the work of AlphaFold2 (Jumper *et al*., 2021), the same group recently formulated a sampling-free “sidechain folding” model that does not rely on using a rotamer library, or energy functions (Xu *et al*., 2022). Along with features used in their previous method, this program utilizes the sidechain density maps derived with another deep learning model (Misiura *et al*., 2021), to train two independent modules: one for predicting the side chain dihedral angles starting from only the backbone and sequence information and the other predicts side chain contact map. Using the contact map as a differentiable scoring function, and the output of the predicted dihedrals as the initial conformation, they deterministically fold the sidechains using a gradient-based method. Although very accurate and entirely a deep learning approach, this method relies heavily on many pre-engineered features taken from the outputs of many different models.

Deep learning methods can discover hidden patterns in the data and complex relationships between interdependent variables. We therefore argue that the sidechain packing problem can be solved without having to rely on pre-engineered features, more directly, using only atomistic coordinate information. For example, a conceptually clear and very intuitive way to solve this problem is to project the protein structure into a 3D voxelated grid and train a 3D CNN model to predict the missing sidechain atoms (Misiura *et al*., 2021). However, a voxelated representation like this would be highly sparse, resulting in inefficient training and poor accuracy. Data augmentation – using multiple orientations for every protein structure data – may also be needed to account for rotation and translation invariance, which adds further to the computational inefficiency of such framework.

Graph Neural Networks (GNN) have gained much popularity in the protein modeling and design community (Sanyal *et al*., 2020; Jing and Xu, 2021; Strokach *et al*., 2020; Gainza *et al*., 2020) due to the fact that graph representation of a protein structure collapses its 3D conformation into a set of nodes and a set of edges defining the relationships between those nodes. Sequence and coordinate information can be embedded as node and edge features and by construction, such representations are invariant upon rotation and translation. A single GNN layer consists of updating all node and edge states in the graph via a neighbourhood aggregation scheme. These layers can be further stacked in order to iteratively refine the state of the graph, resulting in a deep GNN model. The representation vector of a node and the edges is computed and updated by recursively aggregating and transforming node-edge representation vectors of its neighbors defined via an adjacency matrix. The receptive field for the convolution operations in GNN can be limited to, for example in the case of protein molecules, spatially adjacent atoms defining only the local environment. Recently, a GNN based method (Sanyal *et al*., 2020) was developed as an alternative to physics and knowledge-based scoring functions to assess protein structure quality where they use an invariant graph representation of protein.

Using GNN regression models, trained only on crystal PDB data, we develop a computational framework to predict the sidechain geometry that does not rely on a rotamer libary, sampling algorithm, or energy functions. Specifically, we train two sets of models for each sidechain dihedral, each requiring different granularity of the protein structure data, and are employed in-steps to predict the sidechain conformation starting from the protein backbone. The first set of models are used to populate crude conformations of the sidechain which is further iteratively refined using the more enriched second set of models (see Methods). Using non-redundant, high resolution protein structures from the Protein Data Bank (PDB) we use a bidirectional graph representation for each protein. Each protein graph comprises of unique node descriptors specific to amino acid identity and atom-type of a given residue within the protein. The edges are defined using a set of three geometric descriptors that are transformed into standardized features, invariant upon rotation and translation and unique to the protein geometry. We include crystal symmetry mates in the training data to emulate the crystal environment in order to improve model accuracy, as shown in Krivov *et al*. (2009). ZymePackNet uses a GNN architecture based on XENet (Maguire *et al*., 2021) to perform joint message-passing operations on the node and edge attributes.

## 2 Methods

### 2.1 Training Data

The training dataset was obtained using the Pisces server of culled PDB datasets with a sequence similarity *<* 90%, resolution *<* 2.0 Å and a maximum R-factor of 0.25. Only proteins with a maximum of 300 standard amino acids were selected. The structure files obtained from PDB database were processed using GEMMI – an open-source python library for structural biology. Alternate conformations, Het-atoms or structures with missing residues were removed.

Ambiguity in the flip state of Asn and His *χ* and GLN *χ* was addressed using Amber’s reduce -FLIPs tool. This tool, based on the original MolProbity implementation, uses hydrogen bonding network and crystal contact information to identify and selectively correct these side chain dihedrals by 180°. As the model accuracy relies on an accurate depiction of the true environment of an amino acid within the protein, we populated the crystal symmetry mates within 4 Å to the asymmetric unit using CRYST1 records and scale matrices; structures with residue clashes with a crystal neighbor were removed from the dataset. No further optimizations, such as energy minimization of the crystal structure, were performed to avoid biasing or attempting to boost the predictive capabilities of the training model. Approximately 4000 structures were used for training and 1000 structures were withheld for validation.

#### 2.1.1 Protein Graphs

The protein graph representation used here is similar to Sanyal *et al*. (2020) with minor modifications made to work with our prediction framework. The node attributes are chosen as simple categorical variables designating the unique residue name and atom type for every atom in the 20 standard amino acids. The edge attributes are unidirectional and comprise of three types of standardized features, a) the pairwise distance: embedded as 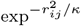, where *κ* is a square of electrostatic cutoff distance – chosen as 10 Å, and *r*_ij_ is the pairwise distance between the two atoms measured in Å, b) a set of three dimensional directional features comprising of the direction of atom *j* from atom *i* using a local coordinate frame constructed using coordinates of the atom *i* and two contiguous bonded atoms (see Supporting Information), to account for apparent anisotropy in relative placement inside the protein and c) a binary input, *b*_ij_, identifying whether the two atoms are covalently bonded. When training the model on a GPU, we found that for larger structures, the entire protein graph could not be processed due to memory limitations. We therefore restrict our models to predict side chain conformations based solely on the local environment of the residue in question, similar in flavor to DLPacker (Misiura *et al*., 2021). We begin by forming a global adjacency matrix such that an edge is created between each atom and its 40 nearest neighbors. The XENet message-passing scheme is specifically designed for symmetric directed graphs with asymmetric edge features (Maguire *et al*., 2021). Therefore, in order to utilize the XENet model, we use a mutual neighbourhood scheme in order to symmetrize our adjacency matrix – if atom *i* is amongst the 40 nearest neighbours of atom *j*, but the converse is not true, we add the missing edge (*i, j*). For each residue, only the subgraph consisting of the residue’s atoms and their neighbors is inputted into the model instance, allowing the model to make graph level predictions. Subgraphs for residues with missing side chain or side chains with low electron density (RSRZ outliers detected using REST API (Mir *et al*., 2018)) were removed from the training data.

### 2.2 Model Architecture

The architecture of a single model in ZymePackNet consists of a node feature embedding layer, followed by two XENet layers (Maguire *et al*., 2021) adopted from the open-source GNN library, Spektral (Grattarola and Alippi, 2021), to perform message-passing operations on the node attributes and edge attributes simultaneously. For the feature stack layer (Maguire *et al*., 2021), pRelu activation was used and for both the outputs of XENet layer, the node and edge attributes, elu activation was used. Each XENet layer was followed by a dropout layer (with a drop out rate of 0.3) and a batchnorm layer. The node features output by the second XENet layer are masked to include only features from a subset of atoms (see Supporting Information) in the residue of interest. These features are passed through an attention sum pooling layer (GlobalAttnSumPool implementation in Spektral(Grattarola and Alippi, 2021) GNN library) which entails a simple attention layer that learns the relative importance of the masked atoms, assigns attention weights to each atom based on importance, and finally computes the weighted average over the entire residue graph. This residue encoding is passed through a single dense layer with a tanh activation, resulting in a 2-dimensional vector representing the sine and cosine of the desired chi angle– sine-cosine representation was chosen over the raw data to account of periodicity.

### 2.3 Training

We train two sets of models which we refer to as partial-context (PC) and full-context (FC), and within each set we train four models, one for each sidechain dihedral angle. While same network architecture is used for each PC and FC model, these models are applied at different stages during prediction and the amount of information inputted as protein graphs are different. Starting from protein backbone, PC models first populates the side chain from *χ* to *χ*, and once all side chains are placed they are iteratively refined using the FC models during prediction. The protein graph used for training PC models on *χ*_i_ is constructed using all backbone atoms and side chain atoms up to and including *χ* for every standard residue in the protein, except for *χ*_1_ where the protein graphs are constructed with protein backbone and C atoms. For FC models however, the graphs are constructed using the backbone atoms and side chain atoms up to and including *χ* for the residue in question, and all backbone and side chain atoms for all other residues in the protein. The atoms needed to construct protein graphs for each stage is listed in the Supporting Information. The mean squared error (MSE) between the network’s predictions and the sine and cosine of the true sidechain dihedral, 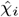, was used to train the network. For symmetric sidechain (Dunbrack Jr and Karplus, 1993) dihedrals *χ*_2_ of ASP, PHE, TYR and *χ*_3_ of GLU, the lower of the two MSE losses pertaining to 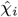 or 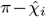 was used. A comparision of the prediction and the ground truth side chain dihedral angles for each of the four models for PC and FC models are shown as scatter plots in the Supporting Information. Each model was trained for 300 epochs with the ADAM optimizer, using an adaptive learning rate and an early stopping criteria, see Supporting Information.

### 2.4 Prediction

The sidechain bond lengths and angles are adopted from the default Amber amino acid parameters library (Case *et al*., 2021). Starting with the protein backbone only, for every residue in the protein, we place dummy sidechains (default Amber coordinates) by aligning their backbone atoms. We then employ the trained PC-*χ*_1_ model on the graph generated using only backbone and the C atoms to predict *χ*_1_’s for all residues in the protein. To apply the predicted *χ*_1_’s, for each residue the dummy sidechain beyond C is twisted about the C -C axis (treating rest of the side chain as rigid object) to match the predicted *χ*. Once all side chains are updated to have the predicted *χ*_1_ we repeat the same process to modify the higher dihedrals using PC-*χ*_2_ through PC-*χ*_4_ ; at each step the predictions are conditioned on the graph generated in the previous step. Using the PC models predicted side chains as initial conformation, we interatively refine our predictions using the FC models. During each iteration, the sidechains are modified sequentially from *χ* to *χ* similar to the PC stage and predictions are conditioned upon the previous graph state. The iteration is stopped when the mean difference in the predictions from two successive iterations is within the desired tolerance, Fig 2.

**Fig. 1.**
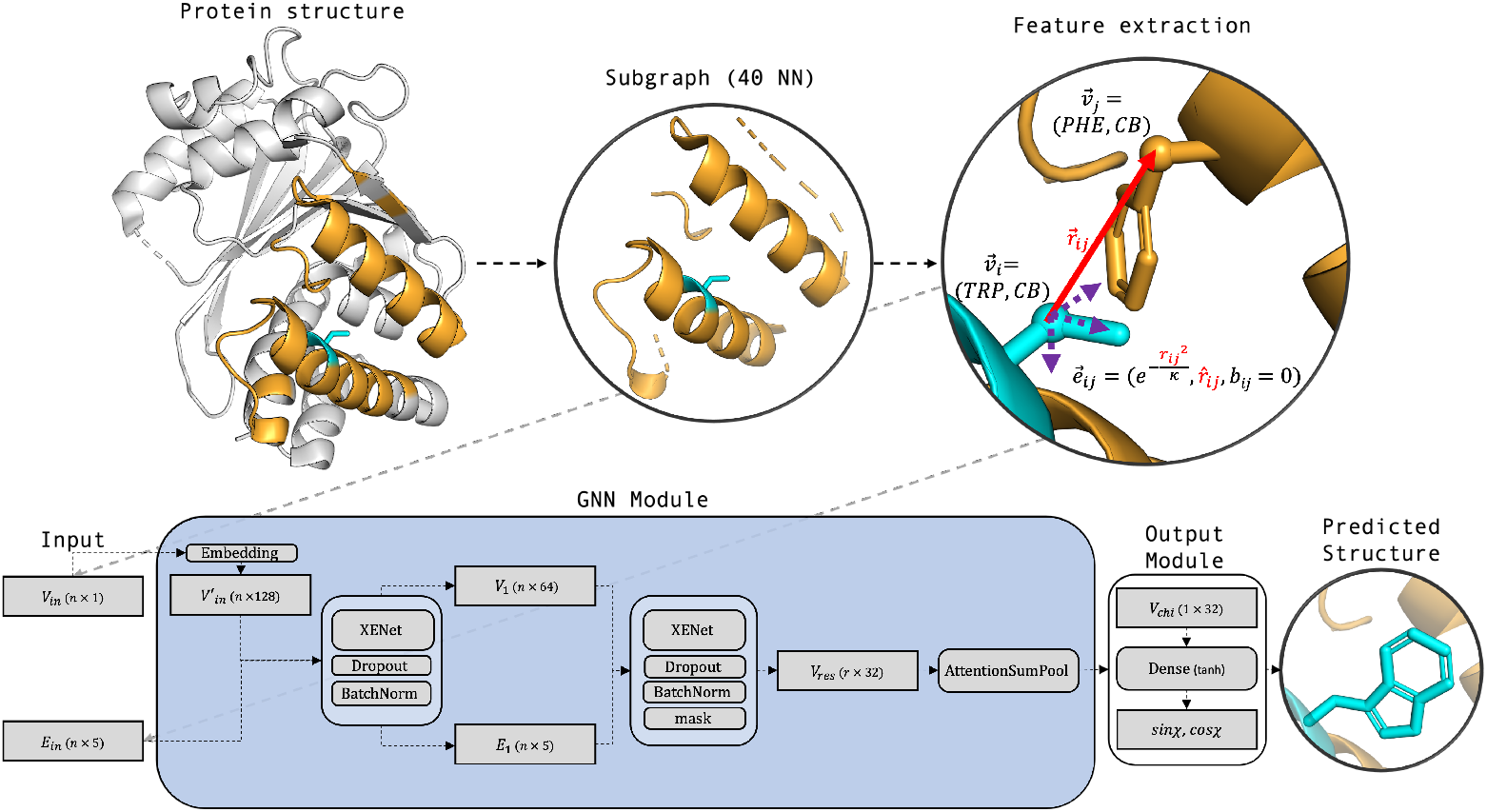
Overview of ZymePackNet. Beginning with the full protein structure, subgraphs for each residue (cyan) are created using the 40 nearest neighbours (orange) of the residue atoms. The node and edge features are extracted from physical (position, distance), and chemical (residue type, atom type, presence of covalent bonding) properties. Only extant atoms are included to construct these subgraphs specific to the query dihedral angle in a specific prediction stage. The GNN module (shown in the lower panel) uses these subgraphs to output the sine and cosine of the desired dihedral angle.

**Fig. 2.**
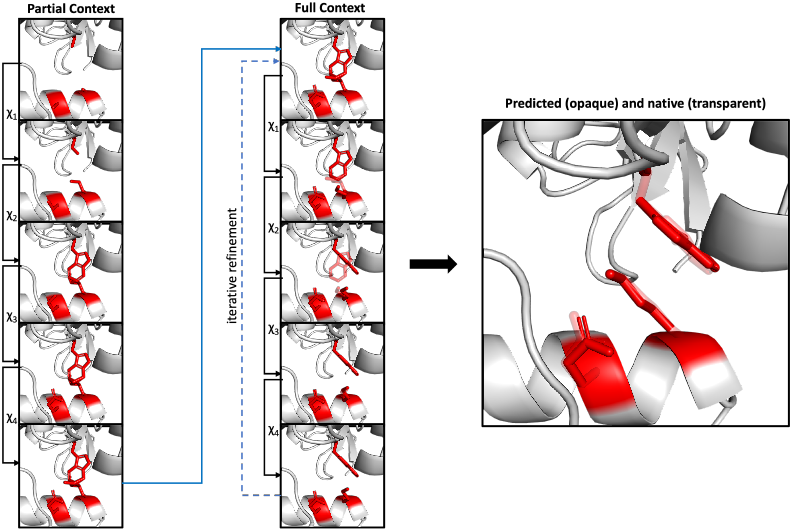
Overview of ZymePackNet prediction protocol. Individually trained Partial Context models are used to progressively grow out the sidechain one *χ*-angle at a time. Afterwards, the Full Context models are applied in sequence in order to refine the model’s predictions. This procedure can be repeated until a desired tolerance in the change between angles of structures from successive iterations is reached.

## 3 Results

### 3.1 Different Refinement Protocols of ZymePackNet

Here we evaluate different refinement protocols and the effect of adding crystal environment for sidechain dihedral angle prediction. Among the three test datasets used in this work, DB379 was the only data set with CRYST1 records and scale matrices required to generate the crystal environment and was therefore chosen for this analysis. We use the same protocol to obtain the symmetry mates as described in the Training Data subsection in Methods. In this construct, the neighbors in the residue subgraphs of the asymmetric unit (AU) may contain atoms from the crystal symmetry mates (SM) if they are among the 40 nearest neighbors to the AU atoms for the residue of interest. While we predict the sidechain dihedrals for residues in the AU only, the same residue level predictions are applied to both AU and mirrored residues in all the crystal SMs. The same criteria was used to populate graphs irrespective of whether crystal contact information was used or not *e*.*g*. PC-*χ*_1_ model only received the graph attributes comprising of backbone and *C*_β_ atoms for all residues.

Apart from assessing the benefit of using the crystal symmetry information, we also experimented with different prediction protocols, see Table 1, – (i) PC: predictions based on PC models without FC refinement, (ii) FC: PC predictions followed by a single FC refinement, and (iii) FC-tol: PC followed by an iterative FC refinement. For Fc-tol, we used an arbitrary convergence criteria *i*.*e*. we stop the FC iterations if the average difference in the predicted dihedrals between two subsequent rounds of FC model is less than the specified tolerance (FC-tol=0.2 rad in Table 1). We used two metrics to quantify the prediction accuracy (i) mean absolute error (MAE) for each of the chi angles in degrees and (ii) ACC, the percentage of the residues in the dataset where all the predicted side chain dihedrals are within 20° of the native structure.

**Table 1.** Performance of adding crystal contact symmetry mates(SM), and different refinement procedures on the DB379 dataset. *χ*_1_ -*χ*_4_ MAE’s are measured in degrees and ACC is measured in %.

| SM | Model | $\chi_1$ | $\chi_2$ | $\chi_3$ | $\chi_4$ | ACC | Time |
| --- | --- | --- | --- | --- | --- | --- | --- |
| ✗ | PC | 17.53 | 24.87 | 44.18 | 54.36 | 61.93 | 1.00 |
| ✗ | FC | 16.10 | 24.03 | 42.44 | 52.52 | 64.02 | 2.02 |
| ✗ | FC-tol | 15.73 | 23.47 | 42.01 | 52.02 | 64.87 | 3.03 |
| ✓ | PC | 16.92 | 24.04 | 43.2 | 54.21 | 63.15 | 2.43 |
| ✓ | FC | 15.26 | 23.08 | 41.52 | 53.38 | 65.42 | 5.34 |
| ✓ | FC-tol | 14.77 | 22.44 | 40.78 | 52.48 | 66.44 | 7.92 |

Evidently, the prediction accuracy can be significantly improved both by adding multiple FC refinement stages, and by adding the crystal environment when making predictions at the expense of added compute. In particular, the FC method significantly outperforms the PC method with twice the compute time, and FC-tol slightly outperforms FC but 1.5 times slower. Adding crystal contacts significantly improves accuracy for all three protocols but amounts to approximately 2.5 higher runtime. Adding crystal environment improves the accuracy on the surface residues, particularly the ones that are involved in making contacts with the symmetry mates, as shown in Fig 3.

**Fig. 3.**
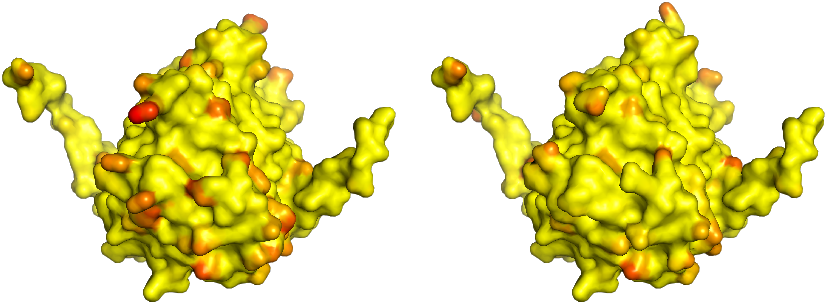
Visualization of improving prediction by adding crystal contact information. Predicted structures using the FC-tol protocol on an example (PDB: 2FW7) from the DB379 dataset, without (left) and with (right) crystal contact information. Atoms close to the native structure are coloured in yellow, while atoms far from the native structure are coloured in red.

### 3.2 ZymePackNet Performance on Three Native Backbone Test Sets

In Table 2 we compare the performance of ZymePackNet with other state-of-the-art methods on three native backbone test sets originally described in Xu et al. (2020): DB379, CASP-FM, and CAMEO-Hard61. For each dataset, we calculate the prediction error over all structures in the respective dataset, in terms of the MAE for each of the four *χ* angles and ACC. Crystal contact information was only available for DB379, and was used to achieve the results reported for each ZymePackNet instance. Numbers for all other methods aside from DLPacker are taken from Xu et al. (2020) For DLPacker, we performed the calculations according to the author’s described procedure.

**Table 2.**
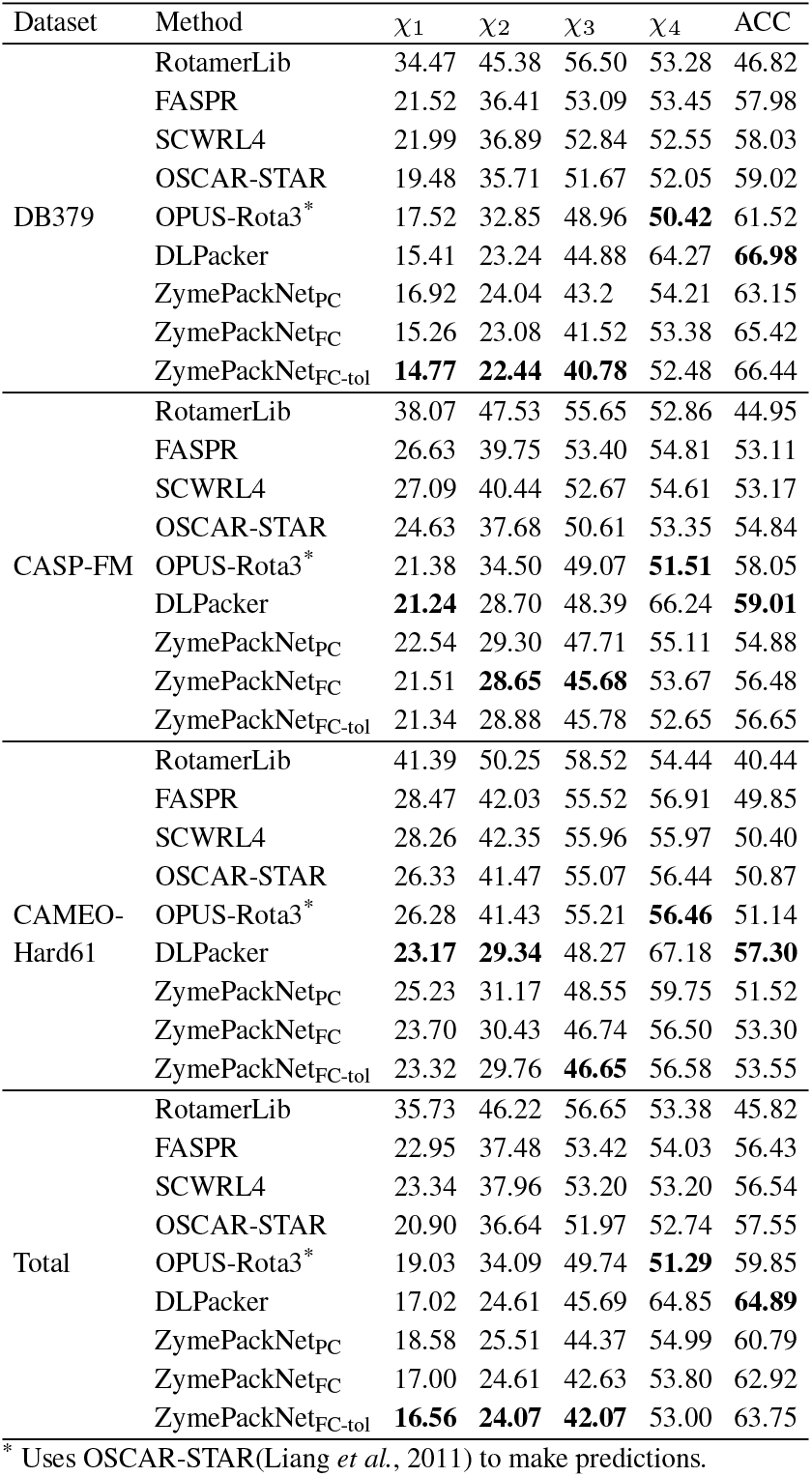
Performance of state-of-the-art sidechain modelling methods on DB379, CASP-FM, and CAMEO-Hard61 datasets. Methods used for comparison with different refinement protocols of ZymepackNet are RotamerLib(Dunbrack Jr, 2002), FASPR(Huang et al., 2020), SCWRL4(Krivov et al., 2009), OSCAR-STAR(Liang et al., 2011), OPUS-Rota3(Xu et al., 2020), OPUS-Rota3(Xu et al., 2020). *χ*_1_ -*χ*_4_ MAE’s are measured in degrees and ACC is measured in %.

From the results above we see the following trend: Without crystal contact information, for *χ*_1_, and *χ*_2_, ZymePackNet, and ZymePackNet achieve marginally worse performance than the best performing alternative – DLPacker. For *χ*_3_, regardless of crystal contact information, ZymePackNet achieves the best performance among available methods. For *χ*_4_, OPUS-ROTA3 achieves the best performance, and when measured according to ACC, DLPacker achieves the best performance. However, when crystal contact information is available (such as for DB379), ZymePackNet achieves better performance than all other methods for *χ, χ*, and *χ*. When compared across all structures in the validation set, ZymePackNet outperforms all alternatives on *χ, χ*, and *χ*_3_, while achieving the third and second-best performance on *χ*_4_, and ACC respectively.

### 3.3 Compute Time Comparison

In order to compare ZymePackNet’s runtime with other methods in the literature, we use the CAMEO-Hard61 dataset as used in Xu et al. (2020). Here we report the running time of ZymePackNet with different refinement protocols, and two other methods FASPR and DLPacker on the same CPU node, Table 3. As FASPR (Huang *et al*., 2020) took the same amount of time (10 s) as reported in Xu et al. (2020), we hypothesize that a similar hardware was used there and therefore the runtime for other methods like SCRWL4, OPUS-Rota3 and OSCAR-STAR reported there should be comparable to what is reported. ZymePackNet_PC_ is the second fastest method reported – slightly faster than SCRWL4. ZymePackNet_FC_ is fifth fastest method, slightly slower than OSCAR-STAR, and moderately faster than OPUS-ROTA3. ZymePackNet is the second slowest method, albeit, significantly faster than the only other pure machine learning method – DLPacker. Overall, ZymePackNet is able to trade off speed at the cost of accuracy.

**Table 3.**
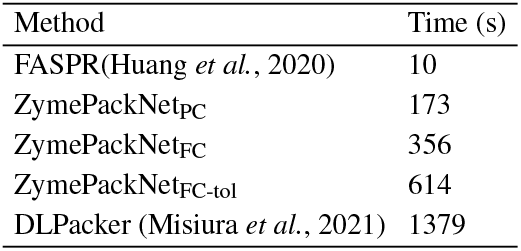
Computing time of sidechain modelling methods on CAMEO-Hard61.

### 3.4 Recovery of *π* Stacking

For sidechain packing algorithms guided by physical energy functions, reconstruction of residue-pair *π* stacking conformations is challenging. The difficulty arises from the fact that there is a range of *π* stacking interactions e.g. *π − π* stacking, *π*-cation, *π*-anion interactions, each requiring a different energy term to account for its energetic effect. Furthermore, these interactions are typically modeled as simplistic two-body interactions, but in reality can manifest due to multi-body interactions. As ZymePackNet uses the local microenvironment, these complicated interactions are predicted without having to explicitly acounting for them in the model, Table 4. In native and repacked structures, these conformations can be detected using a geometric criterion (Brocchieri and Karlin, 1994), given the centroid and *π* plane of each residue contributing to a potential *π* stacking pair (Figure 4). A centroid is either the geometric center of a benzene (PHE, TYR, TRP) or imidozole (HIS) ring, the position of an ARG *C*_*z*_ atom or a LYS *N*_*z*_ atom.

**Table 4.** *π* Stacking recovery metrics on the DB379 dataset.

| Metric | Precision | Recall | F1-score |
| --- | --- | --- | --- |
| Score | 0.82 | 0.86 | 0.84 |

**Fig. 4.**
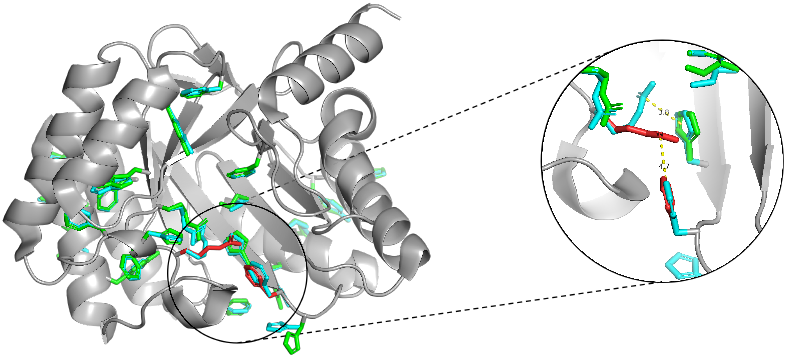
An example (Native 3-dehydroquinase from Salmonella typhi, PDB 1GQN) demonstrating ZymePackNet’s ability to accurately capture *π* stacking. *π* stacking pairs are estimated based on geometric criteria outlined in (Brocchieri and Karlin, 1994). The sidechains involved stacking for the native PDB structure are shown in cyan. ZymePackNet predicted stacks are shown in green for accurately predicted stacks and red for mispredicted.

### 3.5 Interpretability Via Global Attention

ZymePackNet uses an attention pooling layer to intelligently combine the representations of the extant atoms from the residue of interest. In order to better understand which atoms are most important for the network when making predictions, we visualize the attention coefficients produced for each standard amino acid type on an example – PDB 1XDZ from the DB379 dataset. This visualization is provided in Figure 5, where for each PC model, on every standard amino acid, we average and normalize the coefficients produced by the attention layer.

**Fig. 5.**
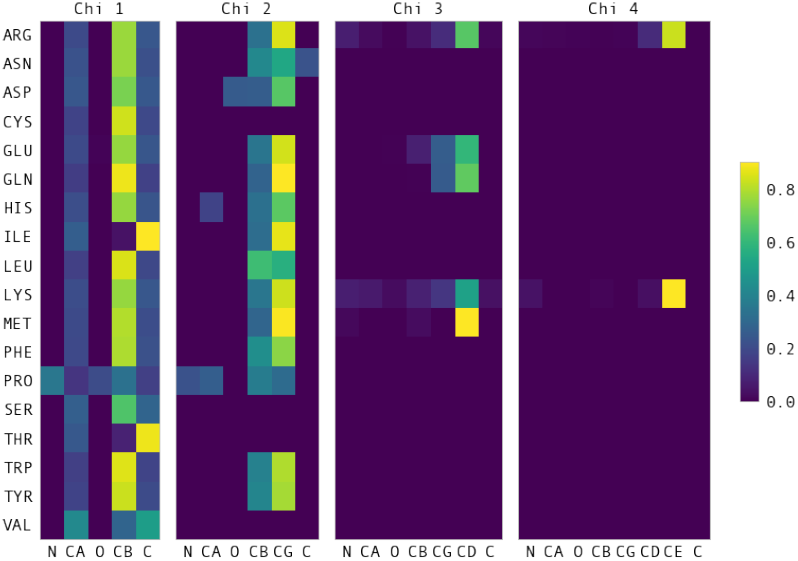
The attention of the final pooling layer visualized for PC-*χ*_1_ *− χ*_4_ over an example (PDB 1XDZ) from the DB379 dataset. The attention coefficients are averaged and normalized for each standard amino acid. Darker (navy blue) square represent areas of low attention, while brighter (yellow) square represent higher areas of attention.

For the most part, the network pays the most attention to atoms anchoring the chi angle being predicted – *C* for *χ, C* for *χ, C* for *χ*, and *C* for *χ*. For specific cases, the network seems to be focusing on other important characteristics. For example, when predicting the first two *χ* angles of Proline, the network pays relatively high attention to the *N* atom, which can be explained by the sidechain of proline forming a pyrrolidine loop connecting the *C*_*α*_ and *N* backbone atoms. Another finding is that when predicting the first *χ* angle of Isoleucine and Threonine, the network pays relatively high attention to the *C* atom, while paying low attention to the *C*_β_ atom. Isoleucine and Threonine are the only two proteinogenic amino acids for which *C*_β_ comprises a second chiral carbon alongside *C*_α_. This might make it more difficult for the network to use the encoding of *C*_β_ in order to make predictions, although does not explain why there is particularly high attention on the backbone *C* atom.

The above findings were fairly consistent across the validation set. Although lending some intuition as to how the network is using the information from different atoms to make the final prediction, it is important to keep in mind that vectors over which attention is performed are higher-order representations of atoms in the original structure. Since two rounds of message-passing have already been performed, individual node vectors encode information about atoms and their local neighbourhoods.

## 4 Discussion

Current protein side chain prediction models focus on developing scoring functions and efficient sampling procedures. In this study, we use a simple, interpretable computational framework using deep learning to demonstrate that side chain dihedral angles can be predicted directly by training on protein structure data.

We train a set of neural network regressor models using only crystal PDB structures, and use these trained models to predict the side chain conformations directly without having to rely on sampling from rotamer library or complex energy functions. The computational framework relies on graph represention of protein structures, where the nodes and edges encode geometric and chemical properties of the structure. Due to graphical nature of the input data, in order to improve prediction accuracy, we also add crystal environment for model training and prediction.

The method entails using two separate modules for prediction, one populates the side chains conditioned upon only protein backbone, while the other refines the initial predictions iteratively with the full context of the amino acid’s local environment. One or both modules can be applied to trade off speed with efficiency. When run in the fastest configuration, i.e. only uses the populating module, ZymePackNet produces sidechain conformations more accurately and faster than SCWRL4 – one of the fastest methods available. At the other end of the spectrum, ZymePackNet can use multiple rounds of iterative refinement along with crystal contact information, producing more accurate results than that of the most accurate protein packing methods. We use an attention based pooling module that allows one to interpret the model by elucidating, via the attention weights, the atoms of varied importance in predicting a particular side chain dihedral.

Traditional side-chain packing methods rely heavily on scoring functions, which are often poorly modeled or heavy approximated for computational tractability. In addition to the well-studied interactions, Van der Waals forces, electrostatics and solvation interactions, and hydrogen bonds, there are also less studied interactions such as *π* stacking that are currently lacking from these methods. Despite not relying on any energy function, we demonstrate that the model can actively recover complex physical interactions within the protein structure such as *π* stacking. Whether other methods are capable of predicting *π* stacks is beyond the scope of this work, traditional energy based models typically ignore this from their scoring functions due to lack of reliable models and computational complexity.

One key step in predicting protein conformation from XRay crystallography and cryo-EM density maps is predicting the side chain conformations. These PDB structures serve as the ‘gold standard’ for developing and validating the protein side chain prediction tools. In practice, the closest rotamer from a discrete rotamer library is assigned that matches the density. Such rotamer libraries are also used by traditional methods to sample within the discrete conformational space. Error introduced by assigning the closest rotamer to build the atomistic models from crystallograhy or EM data are therefore not rectified by such sampling based models. While PDB structures are required to develop a side chain packing method, ZymePackNet eliminates the need for rotamer-library-based sampling. We argue that sample errors originating from assigning discrete rotamer in the structures are “averaged out” during training over a diverse set of structures.

One limitation of ZymePackNet is that the model does not explicitly account for steric clashes. Although we did not notice evidence of clashes more that any other standard packing method such as SCRWL4, one can selectively perform energy minimization within specific regions to eliminate occasional atomic clashes. Alternatively, the training loss function can be modified to penalize inter-atomic clashes, as a future improvement to ZymePackNet.

## Acknowledgements

The authors thank Alejandro Gil Ley, Patrick Farber and Charles Stevens for the discussions and providing feedback in improving the quality of this work.

