## Supplemental Text for "ZymePackNet: rotamer-sampling free graph neural network method for protein sidechain prediction"

#### Supporting Information

Abhishek Mukhopadhyay,*,† Amit Kadan,† Benjamin McMaster,†,‡ J. Liam McWhirter,† and Surjit B. Dixit†

†*Zymeworks Inc, Vancouver, BC, Canada*

‡*University of British Columbia, Vancouver, BC, Canada*

#### Anchor atoms and prediction stage

Table 1: Anchor atoms (A, C) used to compute local reference frame of atom B using according to Eq. 1. Also shown is the last column, the prediction stage where atom B is included in the protein graph.

| Residue | B | A | C | pred_stage |
|---|---|---|---|---|
| all | $N^1$ | $C^{prev2}$ | $CA^{prev2}$ | $1\text{-}\chi_{last}{}^3$ |
| | $CA^1$ | N | $C^{prev2}$ | $1\text{-}\chi_{last}$ |
| | $C^1$ | CA | N | $1\text{-}\chi_{last}$ |

| | O[1] | C | CA | 1-$\chi_{last}$ |
|---|---|---|---|---|
| all and not GLY | CB[4] | CA | N | 1-$\chi_{last}$ |
| ARG | CG | CB | CA | 2-4 |
| | CD | CG | CB | 3-4 |
| | NE | CD | CG | 4 |
| | CZ | NE | CD | N/A[5] |
| | NH1 | CZ | NE | N/A |
| | NH2 | CZ | NE | N/A |
| ASN | CG | CB | CA | 2 |
| | ND2 | CG | CB | N/A |
| | OD1 | CG | CB | N/A |
| ASP | CG | CB | CA | 2 |
| | OD2 | CG | CB | N/A |
| | OD1 | CG | CB | N/A |
| CYS | SG | CB | CA | N/A |
| GLN | CG | CB | CA | 2-3 |
| | CD | CG | CB | 3 |
| | OE1 | CD | CG | N/A |
| | NE2 | CD | CG | N/A |
| GLU | CG | CB | CA | 2-3 |
| | CD | CG | CB | 3 |
| | OE1 | CD | CG | N/A |
| | OE2 | CD | CG | N/A |
| HIS | CG | CB | CA | 2 |
| | ND1 | CG | CB | N/A |
| | CD2 | CG | CB | N/A |
| | CE1 | ND1 | CG | N/A |

| | | | | |
|---|---|---|---|---|
| | NE2 | CD2 | CG | N/A |
| ILE | CG1 | CB | CA | 2 |
| | CG2 | CB | CA | 2 |
| | CD1 | CG1 | CB | N/A |
| LEU | CG | CB | CA | 2 |
| | CD1 | CG | CB | N/A |
| | CD2 | CG | CB | N/A |
| LYS | CG | CB | CA | 2-4 |
| | CD | CG | CB | 3-4 |
| | CE | CD | CG | 4 |
| | NZ | CE | CD | N/A |
| MET | CG | CB | CA | 2-3 |
| | SD | CG | CB | 3 |
| | CE | SD | CG | N/A |
| PHE | CG | CB | CA | 2 |
| | CD1 | CG | CB | N/A |
| | CD2 | CG | CB | N/A |
| | CE1 | CD1 | CG | N/A |
| | CE2 | CD1 | CG | N/A |
| | CZ | CE1 | CD1 | N/A |
| PRO | CG | CB | CA | 2 |
| | CD | CG | CB | N/A |
| SER | OG | CB | CA | N/A |
| TRP | CG | CB | CA | 2 |
| | CD1 | CG | CB | N/A |
| | CD2 | CG | CB | N/A |
| | NE1 | CD1 | CG | N/A |

| | | | | |
|---|---|---|---|---|
| | CE2 | CD2 | CG | N/A |
| | CE3 | CD2 | CG | N/A |
| | CZ2 | CE2 | NE1 | N/A |
| | CZ3 | CE3 | CD2 | N/A |
| | CH2 | CZ2 | CE2 | N/A |
| THR | OG1 | CB | CA | N/A |
| | CG2 | CB | CA | N/A |
| TYR | CG | CB | CA | 2 |
| | CD1 | CG | CB | N/A |
| | CD2 | CG | CB | N/A |
| | CE1 | CD1 | CG | N/A |
| | CE2 | CD2 | CG | N/A |
| | CZ | CE1 | CD1 | N/A |
| | OH | CZ | CE1 | N/A |
| VAL | CG1 | CB | CA | N/A |
| | CG2 | CB | CA | N/A |

For the directional edge features, we use the convention prescribed in Sanyal et al. [1]. The local reference frame $(\hat{e_x}, \hat{e_y}, \hat{e_z})$ for every atom of interest ($B$) is computed using the coordinates of the two preceding contiguous (bonded) atoms $A$,and $C$.

$$\hat{e_z} = \frac{\vec{AB} - \vec{BC}}{\left|\vec{AB} - \vec{BC}\right|}$$

[1] Coordinated taken directly from PDB

[2] $C^{prev}$ and $CA^{prev}$ refers to the preceding (N-term) carbon and alpha carbon

[3] Atom inclusion for building protein graph to the last side chain dihedral, *e.g.* for ARG these atoms are needed for $chi_1$ to $chi_4$, for ASN only for $\chi_1$ and $\chi_2$

[4] Obtained using the backbone atoms and canonical bond and angle relations

[5] Required for populating local environment in FC models

$$
\begin{aligned}
\hat{e_y} &= \frac{\vec{BC} \times \vec{AB}}{\left|\vec{BC} \times \vec{AB}\right|} \\
\hat{e_x} &= \hat{e_z} \times \hat{e_y}
\end{aligned}
\qquad (1)
$$

### Model Parameters and Training

An initial embedding layer converts the categorical node features of unique residue name and atom type to an outer dimension of 128. This is followed by two `XENetConv` layers[2] adopted from the Spektral implementation;[3] `stack_channels` is set to [64, 64], `node_channels` is set to 64, and `edge_channels` is set to 5. The second `XENetConv` layer has `stack_channels` set to [32, 32], `node_channels` set to 32, and `edge_channels` set to 5. The output edge representation from the second `XENetConv` layer is suppressed and omitted from being passed on to the `GlobalAttentionSumPool` layer. For both `XENetConv` layers we set both `node_activation` and `edge_activation` to use the `elu` activation. Each `XENetConv` layer is followed by both a `batchnorm` and `dropout` layer with the dropout rate set to 0.3.

We use an adaptive learning rate initialized to 0.005, which is decreased by a factor 0.2 if the model is unable to reduce the training loss for 5 consecutive iterations. We end training if the validation loss does not improve for 10 consecutive iterations.

### Prediction vs Ground truth

In figures 1-8, using the final trained models, we plot the predicted side-chain dihedral against the PDB data in the validation set. For the symmetric dihedrals, between $\chi$ and $\chi - \pi$ we chose the one pertaining to lower loss, exactly how we describe in the main text.

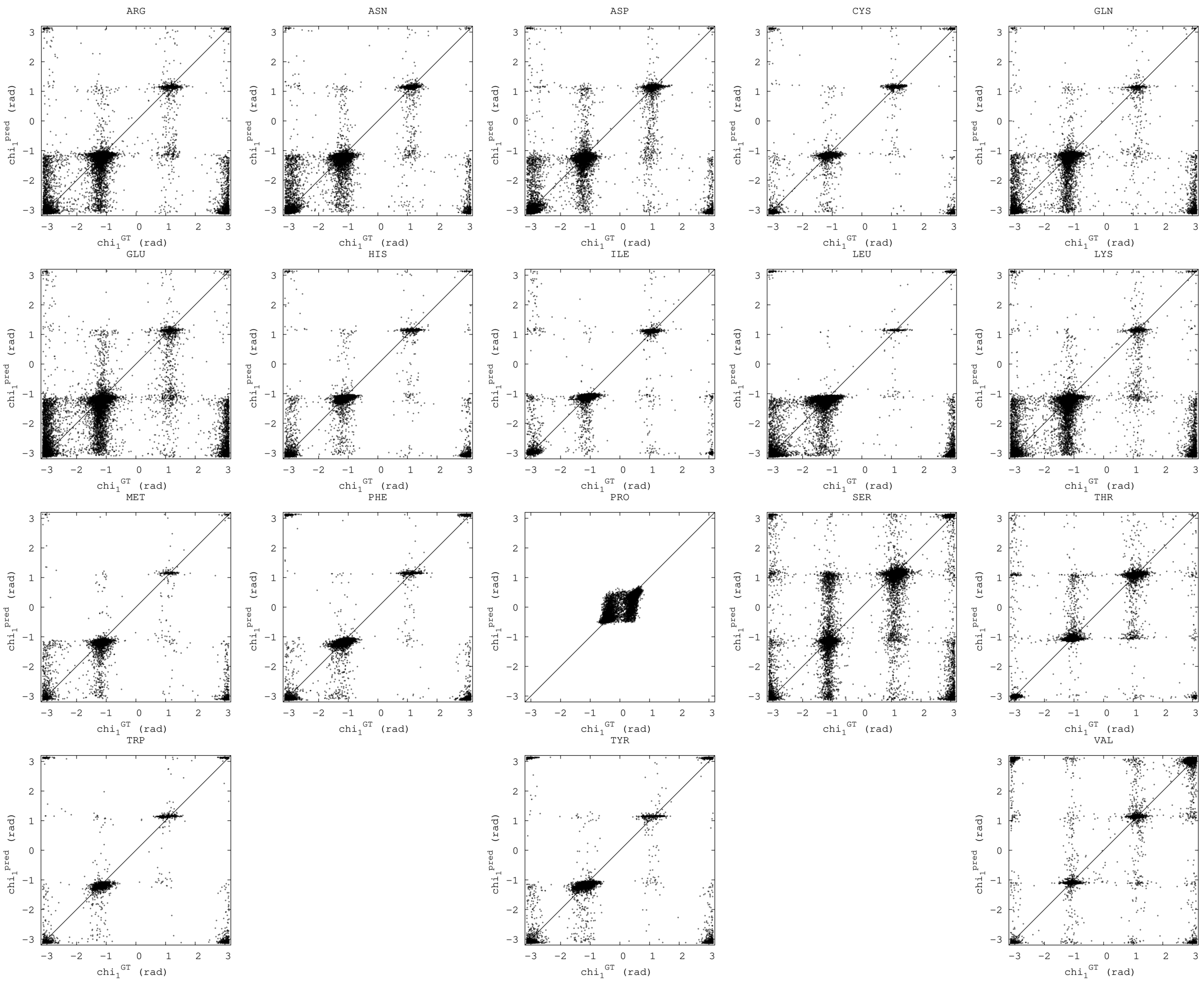


Figure 1: Partial Context $\chi_1$

### Percent accuracy

Apart from the mean average error that is used to compare various side chain packing models, we also use a metric similar to the one in Krivov et al.[4] : percentage of dihedrals that are within 40° of the native structure. For comparison we use ZymePackNet$_{FC-tol}$ model and the results from SCRWL4, see Table 2.

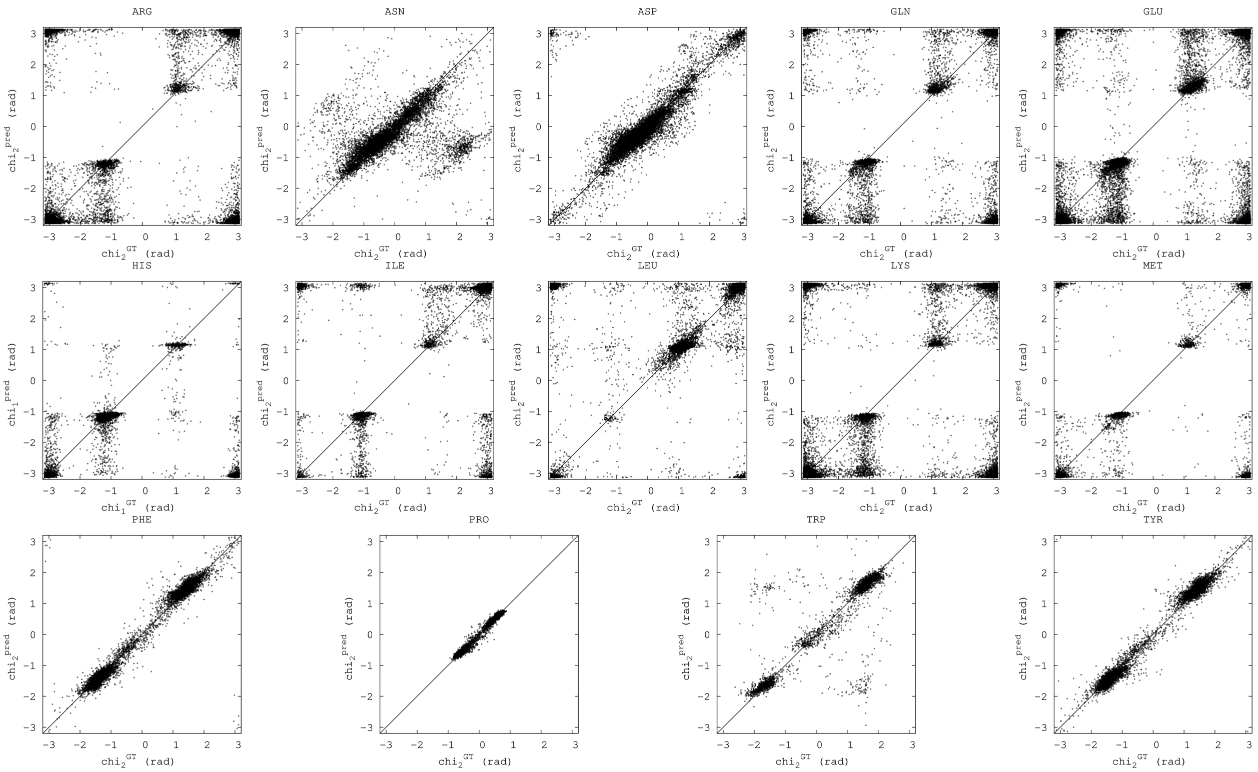


Figure 2: Partial Context $\chi_2$

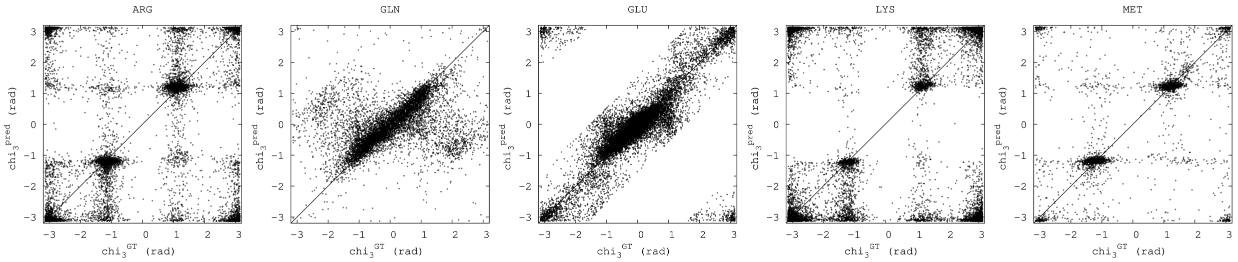


Figure 3: Partial Context $\chi_3$

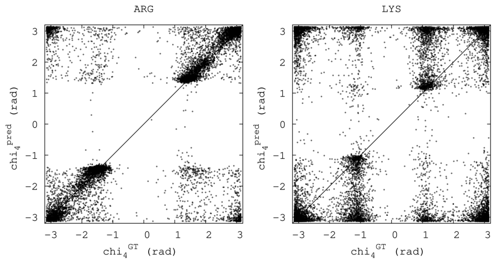


Figure 4: Partial Context $\chi_4$

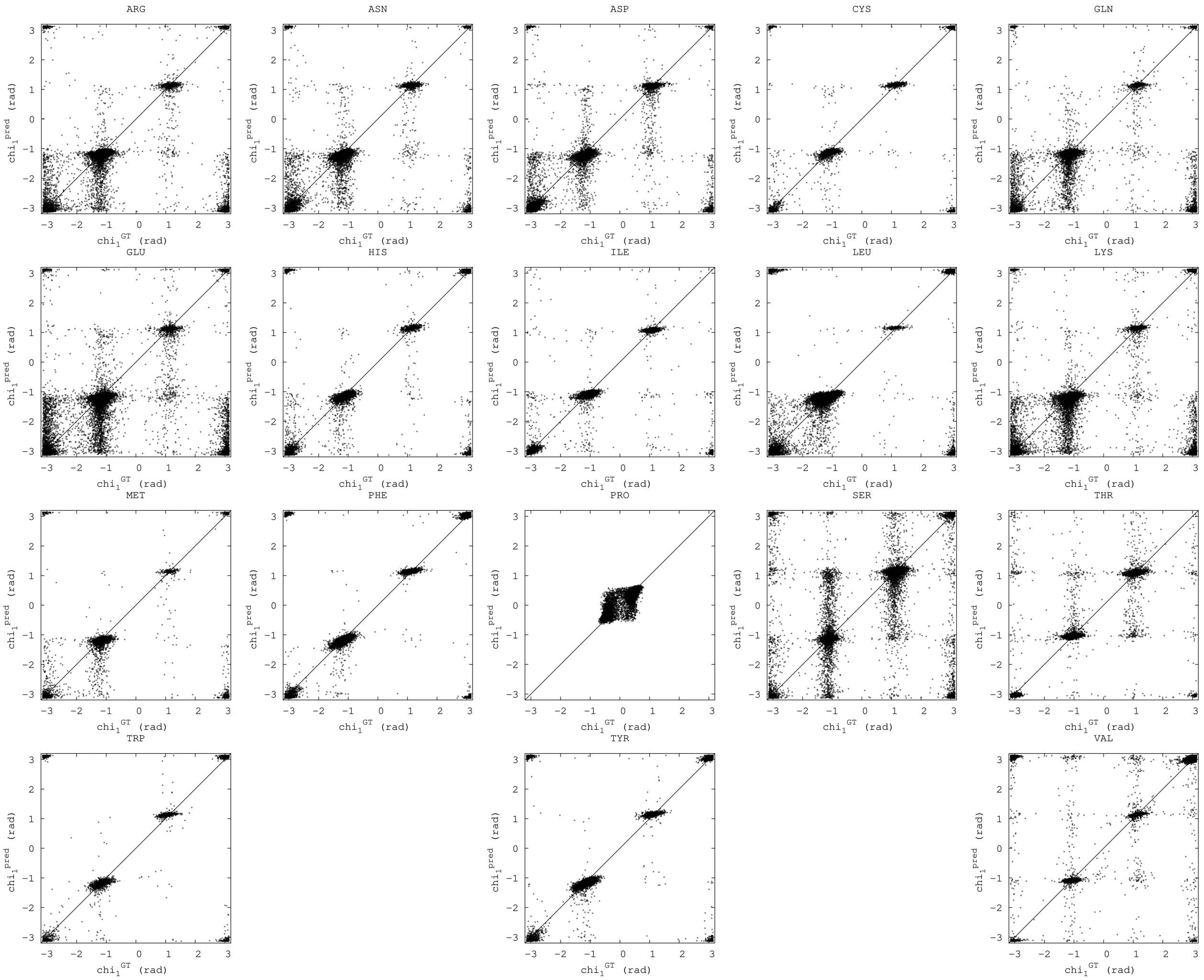


Figure 5: Full Context $\chi_1$

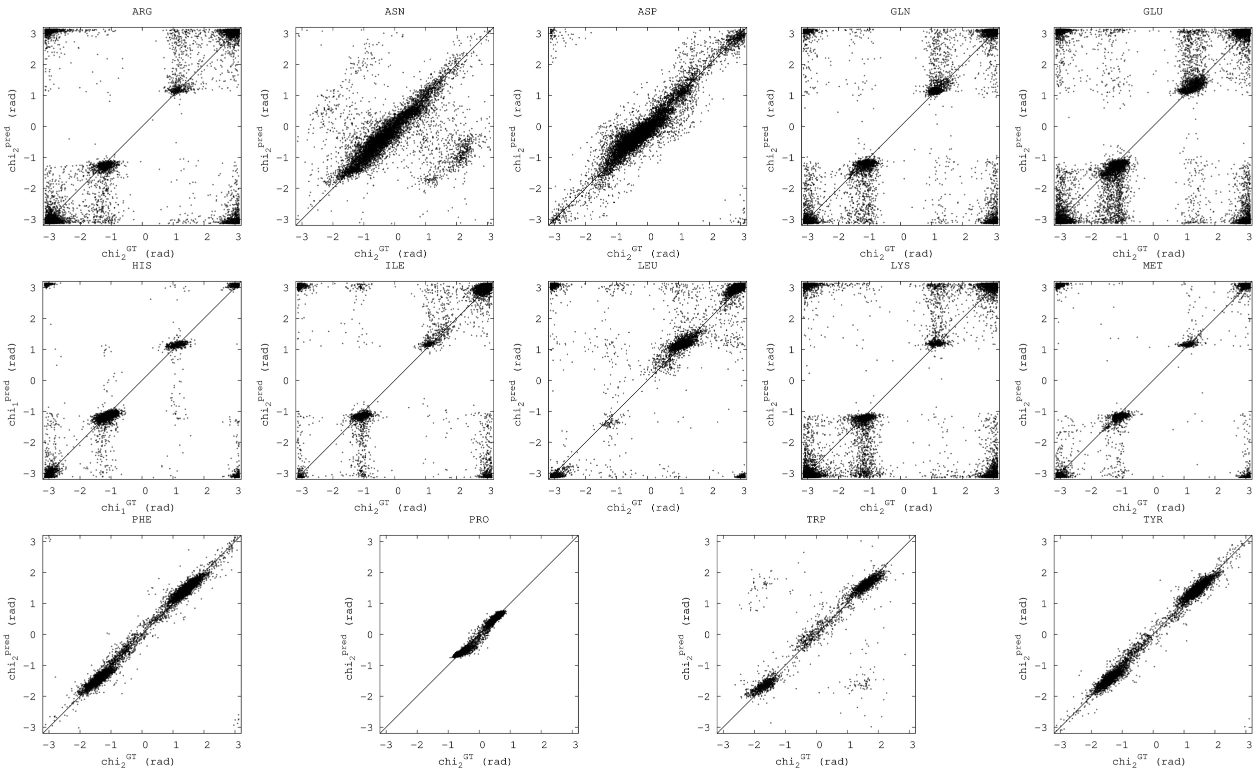


Figure 6: Full Context $\chi_2$

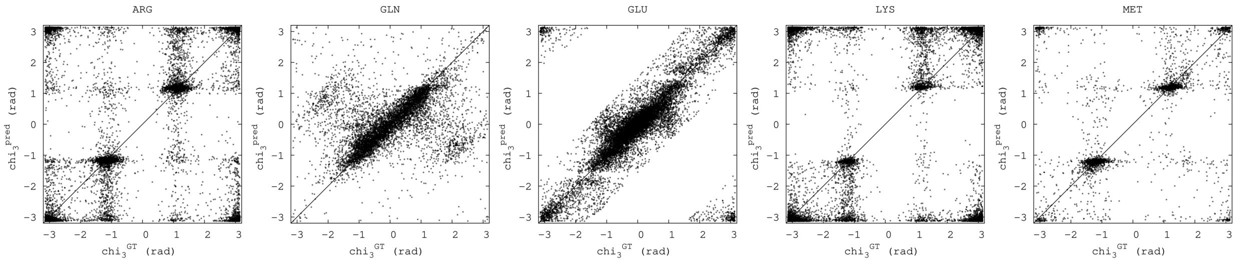


Figure 7: Full Context $\chi_3$

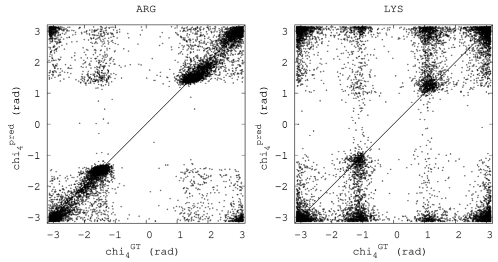


Figure 8: Full Context $\chi_4$

Table 2: Percent accuracy of all side chain dihedrals within 40° of the native structure for ZymePackNet(ZPN) compared with SCRWL4(SC4). ZymePackNet calculations were performed using the FC-tol protocol on the DB379 test set. SCRWL4 values are taken as-is from from the absolute accuracy metric in Table 1 at Krivov et al.[4]

| ResName | chi_1 | | chi_2 | | chi_3 | | chi_3 | |
|---|---|---|---|---|---|---|---|---|
| | SC4 | ZPN | SC4 | ZPN | SC4 | ZPN | SC4 | ZPN |
| ARG | 81.8 | **85.8** | 70.5 | **81.9** | 46.8 | **59.0** | 35.6 | **52.8** |
| ASN | **90.1** | 89.0 | 74.9 | **78.1** | - | - | - | - |
| ASP | 88.8 | **89.7** | 81.8 | **86.8** | - | - | - | - |
| CYS | 92.7 | **95.8** | - | - | - | - | - | - |
| GLN | **84.6** | 83.0 | 67.6 | **74.9** | 46.3 | **56.7** | - | - |
| GLU | 78.3 | **78.9** | 63.8 | **74.6** | 48.2 | **67.5** | - | - |
| HIS | 91.1 | **92.5** | 62.3 | **65.5** | - | - | - | - |
| ILE | **98.6** | 97.2 | **90.9** | 89.3 | - | - | - | - |
| LEU | 95.4 | **96.5** | **91.0** | 89.7 | - | - | - | - |
| LYS | 81.9 | **85.8** | 69.6 | **81.7** | 55.4 | **71.9** | 36.4 | **59.1** |
| MET | 89.0 | **89.4** | 79.0 | **84.2** | 60.9 | **61.2** | - | - |
| PHE | 96.9 | **97.7** | 94.8 | **97.7** | - | - | - | - |
| PRO | 88.2 | **96.9** | 84.7 | **88.8** | - | - | - | - |
| SER | 75.8 | **80.0** | - | - | - | - | - | - |
| THR | 94.0 | **93.8** | - | - | - | - | - | - |
| TRP | 93.0 | **96.1** | 83.0 | **85.9** | - | - | - | - |
| TYR | 95.6 | **97.2** | 93.2 | **97.3** | - | - | - | - |
| VAL | **97.1** | 95.6 | - | - | - | - | - | - |
